# Determinants of dicentric chromosome breakage in *Drosophila*

**DOI:** 10.64898/2026.07.09.737500

**Authors:** Jackson T. Ridges, Hunter J. Hill, James G. Baldwin-Brown, Kent Golic, Nitin Phadnis

**Affiliations:** School of Biological Sciences, University of Utah, Salt Lake City, UT 84112, USA; Institute of Molecular Biotechnology of the Austrian Academy of Sciences (IMBA), Vienna BioCenter (VBC); 1030 Vienna, Austria; Division of Biological and Biomedical Sciences, University of Montana, 32 Campus Drive, Missoula, MT 59812, USA

## Abstract

Eukaryotic genomes often have fragile sites where chromosomes are particularly prone to break. In *Drosophila*, when dicentric ring chromosomes try to segregate, they break at nonrandom hotspots. Here, we precisely map breakage hotspots produced by dicentric ring chromosomes in *Drosophila*. Our study provides three key results about the nature of dicentric chromosome breakage. First, duplications produced by dicentric ring chromosome breakage are surprisingly complex and involve many structural rearrangements, indicating that healing of these breaks is not a simple process. Second, characterization of one particular hotspot showed that new termini all occurred within a single intron of a large testis-expressed gene, suggesting that replication-transcription conflict may be a key determinant of chromosome fragile sites. Third, the new ends are often located near preexisting transposons, suggesting that transposon insertions may contribute to fragility or participate in stabilization of broken ends.

## Introduction

When the attached centromeres of dicentric chromosomes attempt to segregate during cell division, chromosome breakage can occur^1,2^. Previously, we designed a system in *Drosophila melanogaster* that uses sister chromatid exchange in ring chromosomes to generate double-bridge dicentric chromosomes that break during mitotic divisions in the male germline^3,4^ (Figure 1A). When the two bridges break at similar locations and the ends are healed via *de novo* telomere addition, a near-euploid linear-*X* chromosome can be recovered in viable progeny. Our previous cytological studies using polytene chromosomes showed that dicentric chromosome breakage is not random but instead primarily occurs in a limited number of hotspots^4^.

**Figure 1:**
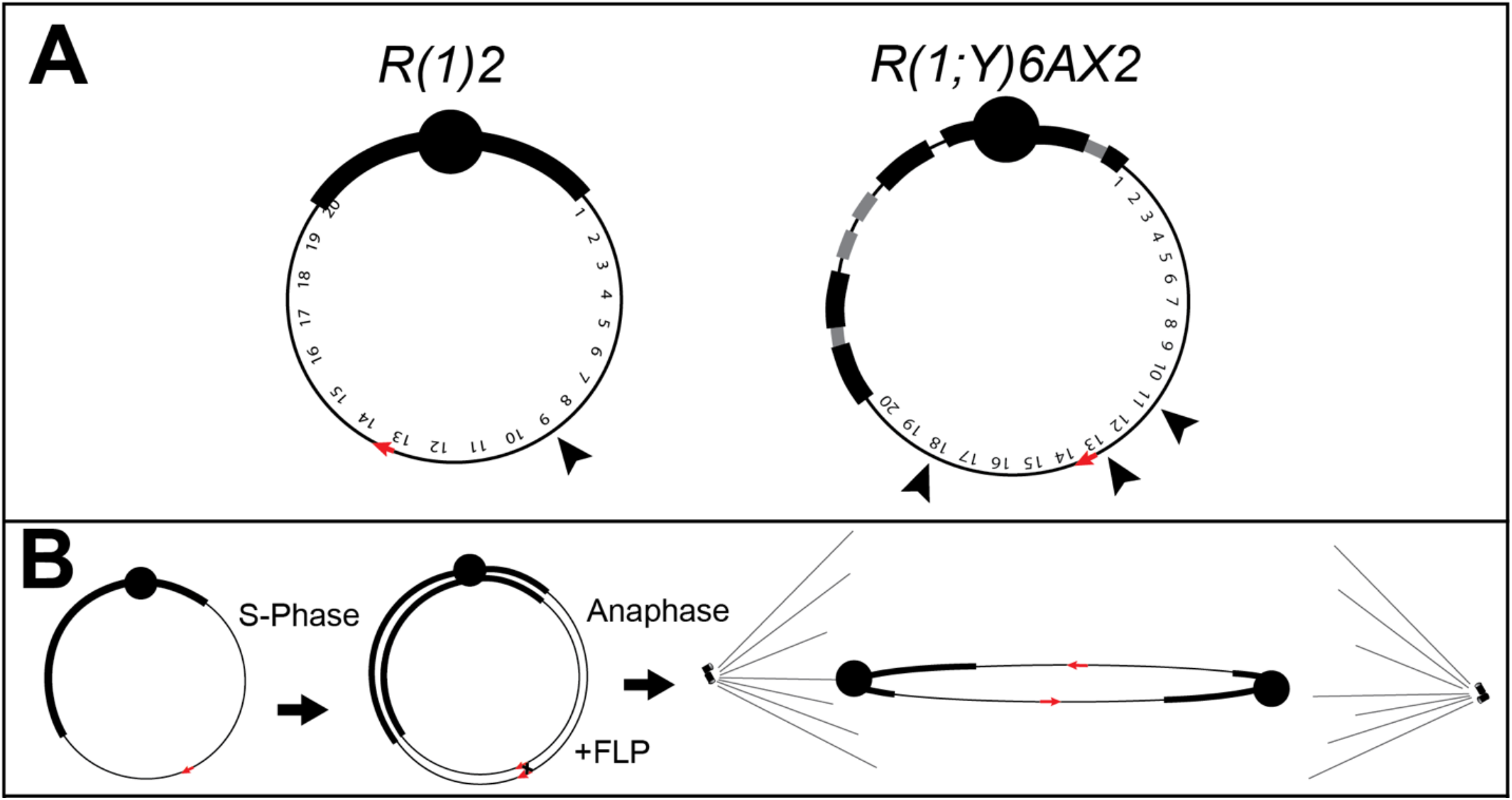
Results of previous studies of broken dicentric chromosomes using our method. **(A)**Previously determined breakage hotspots by cytogenetics of the two ring chromosomes used in this study. **(B)** The strategy used to generate broken and healed linear chromosomes from an initial ring chromosome. FLP mediates sister chromatid exchange at *FRT*s (red arrows) carried on the ring to generate a double-bridge dicentric chromosome which breaks in anaphase in germline mitoses. Broken chromosomes are recovered by mating.

One limitation of our previous work is that we determined breakage location by cytogenetic analysis of linearized ring chromosomes. This leaves open the possibility that breaks tend to occur in the same general chromosomal location, but that at the sequence level, the breaks are diffuse across a broad region. To more precisely identify the ends of these broken-and-healed chromosomes, we sequenced the genomes of more than 50 linearized chromosome lines that had breakpoints in euchromatin. The precise mapping of breakpoints allows us to draw new conclusions about the factors that influence fragility.

A unifying mechanism for chromosome fragility at hotspots remains elusive. Possible mechanisms to explain breakage hotspots in euchromatin include large gene transcripts involved in R-loops, late/under replication, and transposon activity^4^. The most frequent euchromatic breakage hotspot of one of our ring chromosomes occurs in a single gene that encodes a large, testes-expressed transcript. Moreover, several of these independently derived chromosomes have their termini at nearly identical positions. We also found that breakage often occurs at or near preexisting transposons. These results implicate replication-transcription conflict and transposon insertions in determining sites of dicentric breakage and/or stabilization of the broken ends.

## Materials and Methods

### Generation of linear dicentric chromosomes

To determine where dicentric bridges break in *Drosophila melanogaster*, we used linear chromosomes generated in Hill *et al*. 2023^4^ (Figure 1B). Briefly, dicentric formation was induced in the germline of male flies with a ring-*X* chromosome carrying an *FRT*. When FLP recombinase was expressed in the male germline, recombination between sister chromatids at the *FRT* generated a double dicentric bridge. After bridge breakage in anaphase, male-viable linear chromosomes were recovered in the offspring. Since their recovery, these chromosomes have been maintained in stock for periods ranging from many months to several years, typically with attached-*X* females.

### Mitotic cytology

Prior to genomic DNA extraction for sequencing, we verified that chromosomes remained linear by re-examining metaphase chromosomes as previously described^3^. All stocks remained linear except for one stock (*R(1)2#52*) which had re-circularized, likely by intrachromosomal exchange within the large duplication of 6E-8A carried by that chromosome.

### Sequencing of broken chromosomes

DNA was extracted from 15-20 males per independent linear-*X* chromosome using a QIAGEN DNeasy kit. PCR-Free Library preparation and sequencing was done by the University of Utah High-Throughput Genomics (HTG) Shared Resource. 150x150 paired-end sequencing was performed using the Illumina NovaSeq X sequencer. Quality control of sequencing data was done using FastQC. Adaptors and low quality 5’ and 3’ ends were trimmed using Trimmomatic and reads were aligned to the current *D. melanogaster* reference genome (dm6) using bwa mem^5,6^.

### Analysis of breakpoints

We first generated coverage plots for the *X* chromosome of linearized (i.e. opened ring) chromosomes and intact ring chromosomes using the BEDtools genomecov command with the - ibam options^7^. This generated a ‘bedgraph’ file with a coverage value at each coordinate on the *X* chromosome. We then calculated rpkm at each basepair coordinate for the broken and unbroken ring chromosomes. Then, we divided the broken ring rpkm by the unbroken ring rpkm to get a normalized value at each position. We then calculated sliding window averages of rpkm ratios with a window size of 10 kb and a step of 1 kb. Resulting coverage files were visualized using IGV^8^.

To search for transposon incorporation at the end of linear chromosomes, we found the mates of complete reads at the edge of deletions that face the deletion. We then used BLASTn to compare the sequence of mates to a library of *D. melanogaster* transposon sequences^9^. To define hits, we took the best hit and used a threshold of e-value < 10^-5^. In practice, all our best hits were much lower than this threshold.

### Permutation test for transposon proximity

To perform the permutation test, we first recorded the exact basepair of each breakpoint from linear chromosomes that carry deletions (32 in total). We then accessed the existing gff file containing transposon insertions in the dm6 reference genome from FlyBase^9^. We used our program ‘permuvals’ to randomly permute 32 breakpoints 10,000 times and determine how many permuted breakpoints are within 500 bp of an annotated transposon insertion (https://github.com/jgbaldwinbrown/permuvals). The location of transposon insertions was constrained throughout the permutations. We then fit a poisson distribution to our permuted frequencies of breakpoints adjacent to transposon insertions, which had a *mu* of 2.254. We then calculated the likelihood of observing at least 15 breakpoints (our observed value) within 500 bp of transposon insertions based on this distribution.

## Results

### Complex rearrangements of duplicated regions produced by breakage

To precisely map the endpoints of the linearized chromosomes, we sequenced the genomes of males with linear *X* chromosomes previously generated through our screen. Asymmetric breakage of the two bridges of the dicentric chromosome should produce one linear chromosome with a duplication and its complement with a deletion. For euchromatic breakage events, we expected to primarily recover duplications because deletions are more likely to be lethal in males. In total, we sequenced 56 independently linearized *X* chromosomes from two original ring chromosomes (*R(1;Y)6AX2* and *R(1)2*). We first normalized the coverage of broken rings to that of males with the corresponding unbroken ring chromosome. Since we previously observed that breakage events often result in duplications on the *X* chromosome^4^, we determined regions of the *X* chromosome that show doubled coverage in males. The ends of the linearized chromosomes may then be inferred by determining the boundaries of duplications (Figure 2A). Of these 56 linear chromosomes, 28 carried duplications around the breakpoints, 16 carried small euchromatic deletions that are viable in males, and the remaining 12 had no obvious deletions or duplications, likely because the duplication or deletion is too small to be detected (i.e., both new ends are at nearly the same location) or because chromosome ends are in a region that aligns poorly to the dm6 reference genome.

**Figure 2:**
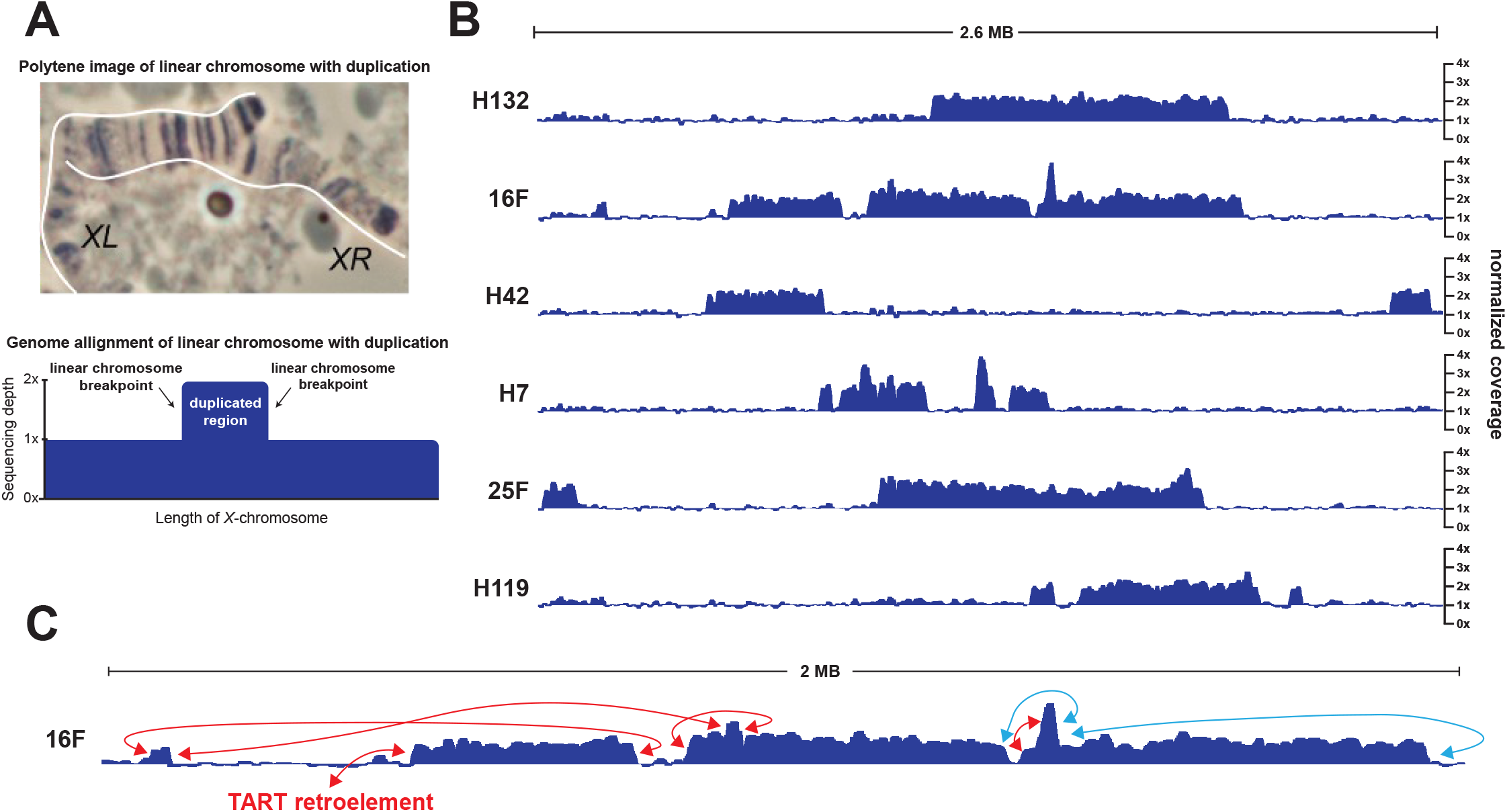
Duplications produced by dicentric chromosome breakage. **(A)** (Top) Polytene image of a broken chromosome that carries a long duplication. The duplicated terminal regions pairs apart from a small segment at one terminus. (Bottom) Schematic of how a simple duplication produced by a breakage event should look when aligned to the *D. melanogaster* reference genome. Edges of the duplicated region should represent the edges of the linear chromosome, and the initial breakpoints. **(B)** Normalized coverage plots showing the relative coverage of five linear chromosomes compared to the unbroken ring (see Materials and Methods). All five lines show large, duplicated regions. A 2.6 MB region of the *X*-chromosome is shown that overlaps the previously described breakage hotspot at polytene band 11. The Y axis ranges from 0x to 4x of relative coverage for each plot. **(C)** Location of read pairs for reads at the edge of duplicated regions. Blue lines indicate paired reads that sequenced the same DNA strand of the reference genome. This analysis suggests that duplications have complex chromosomal rearrangements.

Nearly all chromosomes carrying duplications (25/28) were predicted by polytene cytology. Regions of increased coverage were immediately apparent, and matched the ends observed in polytene cytology, with the largest duplication covering ∼1.9 megabases (Table 1). However, sequence analysis revealed hidden structural variation that was not detectable by cytology. Some duplications were surprisingly complex and appear to have been shaped by multiple events beyond simple breakage and healing, manifesting in alignments as several distinct duplications and as regions with more than 2x coverage (Figure 2B). To attempt to disentangle the structural variations at duplications, we focused on discordant reads at the edges of duplicated regions. Many lines showed complex patterns: *e*.*g*., paired reads that were unexpectedly distant from each other suggesting a duplication or transposition of genomic DNA (1st red arrow in Figure 2C); or paired reads that sequenced the same DNA strand of the reference genome, indicating an inversion (blue arrows, Figure 2C). In some cases, the polytene chromosome cytology was suggestive of this complexity — for instance, Figure 2A shows a duplication-bearing linear chromosome in which one chromosome end is not fully paired and seemingly not homologous to its partner. This complexity prohibited precise identification of the ends of these chromosomes. It is worth noting that we found no instances of autosomal DNA duplication in these genomes, indicating that only *X* chromosome material is found at the ends of the linearized chromosomes.

**Table 1:** Broken ring chromosomes with duplications.

| **Broken ring** | **Initial ring chromosome** | **Polytene location of break** | **Approximate span of duplication/s** | **Approximate number of duplicated segments** |
| --- | --- | --- | --- | --- |
| **H135** | R(1;Y)6AX2 | 4C-4D | 260 KB | 1 |
| **OR31** | R(1;2) | 5E-6F | 870 KB | 2 |
| **OR96** | R(1;2) | 6A-6B | 75 KB | 2 |
| **OR36** | R(1;2) | 7A-7B | 340 KB | 3 |
| **OR19** | R(1;2) | 7B-8C | 1.4 MB | 2 |
| **OR107** | R(1;2) | 7D-8B | 1.1 MB | 2 |
| **H105** | R(1;Y)6AX2 | 7D-8D | 230 KB | 1 |
| **OR84** | R(1;2) | 7D-8E | 1.7 MB | 1 |
| **OR60** | R(1;2) | 7D-8E | 1.5 MB | 3 |
| **OR92** | R(1;2) | 8F-10C | 1.8 MB | 1 |
| **H66** | R(1;Y)6AX2 | 9E-10E | 500 KB | 3 |
| **H123** | R(1;Y)6AX2 | 10B-10F | 190 KB | 1 |
| **H42** | R(1;Y)6AX2 | 10D-11A | 460 KB | 2 |
| **16F** | R(1;Y)6AX2 | 10E-11D | 1.5 MB | 4 |
| **H7** | R(1;Y)6AX2 | 11A | 700 KB | 4 |
| **H94** | R(1;Y)6AX2 | 11A | 560 KB | 1 |
| **25F** | R(1;Y)6AX2 | 11A-11D | 1.9 MB | 2 |
| **H132** | R(1;Y)6AX2 | 11A-11D | 860 KB | 1 |
| **H119** | R(1;Y)6AX2 | 11B-11E | 800 KB | 3 |
| **H24** | R(1;Y)6AX2 | 12D-13A | 880 KB | 2 |
| **H102** | R(1;Y)6AX2 | 12E-13D | 150 KB | 2 |
| **H120** | R(1;Y)6AX2 | 12E-12F | 400 KB | 1 |
| **2F** | R(1;Y)6AX2 | 13A | 230 KB | 1 |
| **HMF11** | R(1;Y)6AX2 | 13E-13F | 75 KB | 1 |
| **H104** | R(1;Y)6AX2 | 13F-14B | 480 KB | 2 |
| **H91** | R(1;Y)6AX2 | 14B-15D | 980 KB | 4 |
| **H8** | R(1;Y)6AX2 | 15A-15F | 550 KB | 1 |
| **HMF24** | R(1;Y)6AX2 | 17C-17F | 410 KB | 2 |

### A breakage hotspot that generated only deletions

We next turned our attention to the 16 lines that carried deletions and found that precise endpoints of deletions could be more readily identified (Table 2; example shown in Figure 3A). We therefore focused the rest of this investigation on linearized chromosomes with deletions.

**Table 2:** Broken ring chromosomes with deletions. We note that polytene analysis placed H34 at a slightly different location than the other region 18 breakpoints, while the sequence analysis shows that all region 18 breakpoints overlap. This most likely reflects the occasional difficulty of accurate polytene analysis.

| **Broken ring** | **Initial ring chromosome** | **Polytene location of break** | **Approximate span of deletion** |
| --- | --- | --- | --- |
| **OR106** | R(1;2) | 5F | 8.5 KB |
| **OR63** | R(1;2) | 6A | 2.8 KB |
| **H93** | R(1;Y)6AX2 | 6F | 13.5 KB |
| **H129** | R(1;Y)6AX2 | 7B | 24 KB |
| **OR69** | R(1;2) | 9A | 9.2 KB |
| **OR110** | R(1;2) | 9A | 27.8 KB |
| **OR93** | R(1;2) | 9A-9B | 1.9 KB |
| **H107** | R(1;Y)6AX2 | 12A-12B | 16.1 KB |
| **OR90** | R(1;2) | 13A | 2 KB |
| **OR13** | R(1;2) | 13C | 15 KB |
| **H6** | R(1;Y)6AX2 | 17D | 8 KB |
| **H125** | R(1;Y)6AX2 | 18B-18D | 77.4 KB |
| **H97** | R(1;Y)6AX2 | 18C | 9.7 KB |
| **H103** | R(1;Y)6AX2 | 18C | 20.6 KB |
| **HMF6** | R(1;Y)6AX2 | 18C-18D | 10 KB |
| **H34** | R(1;Y)6AX2 | 18F-19A | 8 KB |

**Figure 3:**
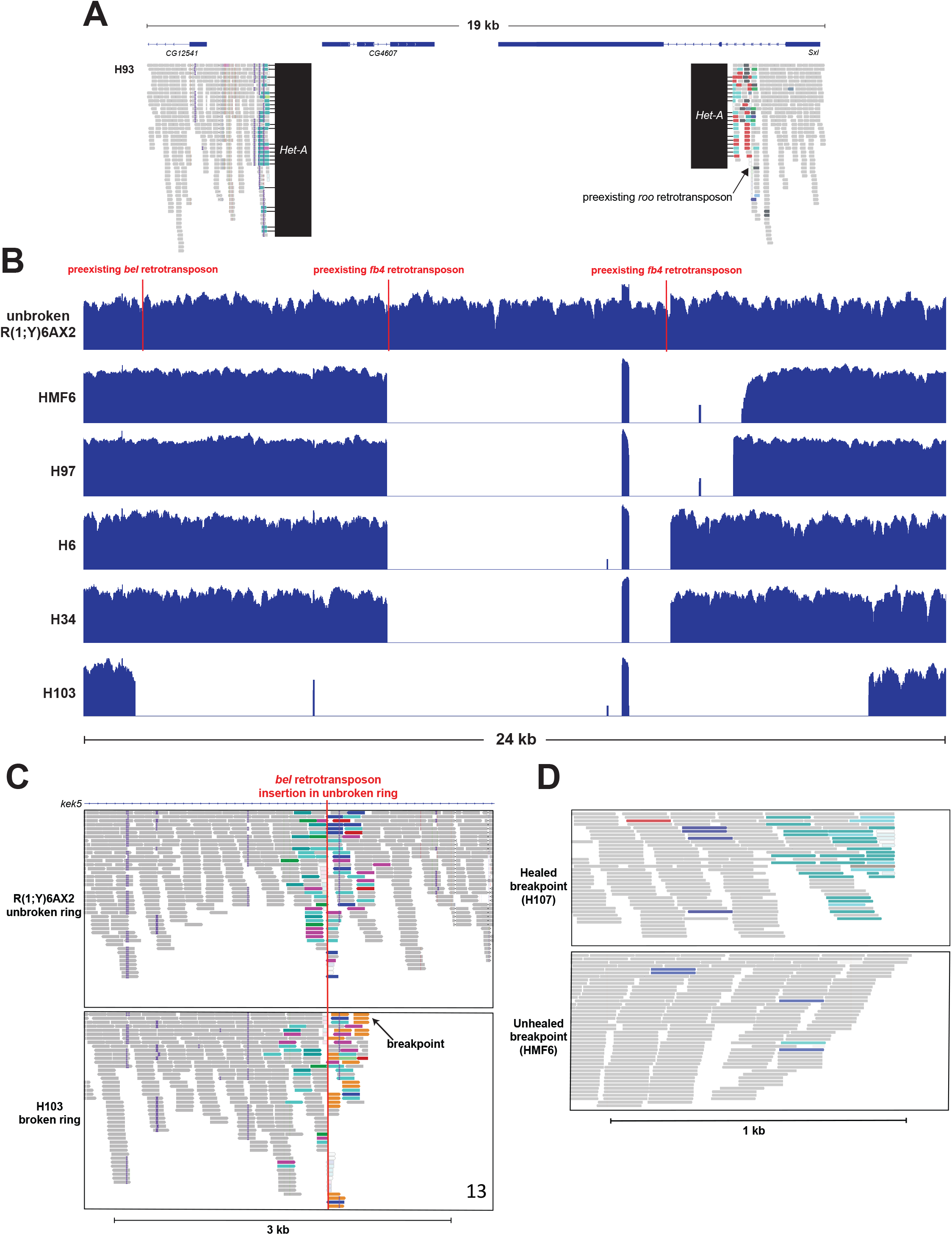
Deletions produced by dicentric chromosome breakage. **(A)** Alignment of the deletion produced by breakage in line H93. The deletion has healed through addition of the telomeric retrotransposon *Het-A* at both breakpoints. The right breakpoint occurred at the location of a preexisting *roo* transposon in the unbroken ring chromosome. Reads that are any color apart from grey or white have discordant read pairs that map to *roo* insertions elsewhere in the genome. **(B)**Unnormalized coverage plots of several independent linear chromosomes that carry overlapping deletions at the breakage hotspot in polytene band 18. These deletions are all in the first intron of *kek5*. Three retrotransposon insertions are shown that are present in the unbroken ring chromosome. Note that the small regions of coverage in the middle of deletions are repetitive regions with read fragments that map nonspecifically to this region. **(C)** Alignment of a breakage site in line H103 that occurred at a preexisting *bel* transposon insertion in the unbroken ring chromosome. Reads that are any color apart from grey or white have discordant read pairs that map to transposon insertions elsewhere in the genome. **(D)** Alignments of two breakpoints at the edge of duplications in lines H107 and HMF6. In H107, the breakpoint has healed through addition of *Het-A*, a telomeric retrotransposon, which was determined by the sequence of the paired read of reads at the breakpoint. In line HMF6, the breakpoint has remained unhealed, and coverage gradually decays as the alignment approaches the breakpoint. Reads that are any color apart from grey or white have discordant read pairs that map to transposon insertions elsewhere in the genome.

There was a strong breakage hotspot in ring chromosome *R(1;Y)6AX2* in polytene division 18. We discovered that breakage events at this hotspot resulted in deletions rather than duplications: specifically, all six linear chromosomes that we sequenced with breakpoints at this hotspot carried deletions in the first intron of *kekkon-5* (*kek5*). These deletions overlap and range in size from 10 kb to 80 kb (Figure 3B, largest deletion not shown). Three deletions at the breakage hotspot of polytene division 9A in *R(1)2* were within a 200 kb genomic segment but were non-overlapping and occur within different genes (not shown), emphasizing the unique nature of the division 18 hotspot.

Aspects of the breakage hotspot in 18 are consistent with multiple previously proposed hypotheses to describe chromosome fragile sites^4^. *kek5* is an abnormally large gene in *Drosophila* (∼120 kb), and publicly available RNA-seq datasets show that it has moderate expression in the male germline^9^. This leaves open the possibility that either R-loops formed by large transcripts or replication-transcription conflict may underly this hotspot^10^. Additionally, we found that there are an unusually high number of transposon insertions within this intron of *kek5* in the unbroken ring chromosome (6 transposon insertions in ∼ 110 kb) (3 shown in Figure 3B). This raises the possibility that this site is prone to chromosome breakage because of the presence of transposons.

### Enrichment of breakpoints near transposons

We next wondered whether proximity to transposon insertions is a common theme among breakage hotspots. We determined the distance from the nearest transposon insertion for all breakpoints that resulted in deletions and found that 15 out of 32 total chromosome ends occurred within 500 bp of a preexisting transposon insertion (∼47%, example shown in Figure 3C). For many breakpoints that produced deletions, discordant reads have pairs that map directly to transposons, suggesting that the chromosome end is at or within an existing transposon. To determine if the frequency of breakpoints in or adjacent to transposons is higher than expected by chance, we performed a permutation test. We found that our observed frequency of chromosome ends within 500 bp of a transposon insertion is significantly higher than expected from the distribution of our permuted sets of breakpoints (see materials and methods, 10,000 permutations, *p* < 0.0001).

### Addition of telomeric retrotransposons

We next investigated the ends of linear chromosomes to determine whether they have acquired telomeric retrotransposons. *Drosophila melanogaster* lacks telomerase and telomeres are maintained by domestication of three retrotransposons: *Het-A, TART*, and *TAHRE* (HTT). These retrotransposons repeatedly attach themselves to the ends of linear chromosomes to maintain chromosome length and resolve the end replication problem^11^. The ends of linear chromosomes that we recovered must possess the multi-protein cap, called terminin^12^, that prevents the cell from detecting the chromosome end as a double strand break. However, assembly of the terminin cap does not require a particular DNA sequence at the end of the chromosome and such ends do not, at least in the short term, require telomeric retrotransposons^13^. Previous work has shown that broken chromosome ends can occasionally acquire the HTT retrotransposons^14–16^, and some of these linear chromosomes have done so (7/32; ∼22%). Both termini of H93 show *Het-A* addition (Figure 3A), while the right side terminus of H97 (as diagramed in Figure 3B) and the left side of H103 have both received *TAHRE* additions. This is likely an underestimate of the total number of breakpoints that are healed by telomeric retrotransposon integration, as breakpoints that occurred within existing retrotransposons can’t be analyzed by this method. We also noticed that some of the breakpoints seem to remain unhealed (11/32; ∼34%), with a gradual decrease in read depth approaching the breakpoint, no doubt owing to incomplete replication of chromosome ends (*e*.*g*., HMF6, Figure 3D). Within such males there were sometimes a few reads that terminated in HTTs, suggesting somatic addition of HTTs or that DNA was prepared from a mixed population with a few males carrying HTT additions. While some linear chromosomes have clearly healed through *de novo* addition of telomeric retrotransposons, others continue to suffer from degradation owing to incomplete replication.

## Discussion

Under conditions of replication stress, such as starvation for nucleotides or their precursors, cells experience incomplete replication manifesting as constrictions in metaphase chromosomes and repeated breakage at loci known as common fragile sites (CFS)^17^. In humans, progress has been made towards understanding the disproportionate number of breaks that occur at common fragile sites. It is thought that these regions break more often primarily because they are unusually slow replicating or located far from replication origins^18^. The chromosomal breaks we study here are produced through a different method – tension-induced breakage caused by two attached centromeres segregating to separate daughter cells. We have previously shown that these chromosomes tend to repeatedly break in particular cytological regions. We undertook whole genome sequencing of flies with the recovered broken chromosomes to more precisely assess the locations of the chromosome termini and to deduce the causes of breakage hotspots.

A surprising result was the unexpected complexity of duplications produced by breakage. There is no obvious *a priori* reason to expect duplications to be more complex than deletions, but they frequently involved complex chromosomal rearrangements and multiple distinct duplicated segments. Although this complexity impeded our ability to precisely determine chromosome ends, it raises questions about the nature of breakage events and the stability of broken chromosomes. Simple duplications could be generated by asymmetric breakage of the twin dicentric bridges and healing of broken ends, but more complex duplications require additional steps. For instance, if the two ends of a broken chromosome with a duplication fused prior to S phase, a ring with a tandem duplication would be generated. FLP recombinase might then again generate a dicentric subject to breakage in the next division and further rearrangement. Moreover, if a broken chromosome replicated without healing, fusion of sister chromatids at the same chromosome end would generate a dicentric, and breakage of the resulting bridge in the next division could generate an end with an inverted duplication. The combination of these two types of events, possibly recurring over several mitotic cycles, could generate complex rearrangements. Rearrangements such as these are characteristic of some human aneuploidies, such as 8p syndrome^19^, and are known as INV-DUP-DEL rearrangements^20^. Recombination and chromosome breakage-based mechanisms for their formation have been discussed extensively^20–23^. Alternatively, break-induced replication (BIR) combined with template switching is capable of producing complex rearrangements in a single cell cycle^23–25^.

Since several fly generations elapsed between recovery of a linearized chromosome and extraction of DNA for sequencing. It is unknown whether the observed complexity was generated immediately after breakage or accumulated later. Perhaps duplications were initially simple but later lost fragments within the duplication. If the dosage imbalance produced by duplicated genes is deleterious, deletions that restore correct copy number may have a selective advantage. Although linearized chromosomes were generally kept so that they passed only through males, where there is no meiotic recombination, it is possible that mitotic recombination events occurred and altered the initial structure of the broken ends. In fact, the identification of a linear chromosome that initially carried a long duplication but had recircularized shows that this can occur (see Materials and Methods). Long-read sequencing of newly linearized chromosomes may clarify whether these complex rearrangements are produced during breakage or accumulate later and provide insight into the mechanism(s) by which they occur.

Though the majority of linearized chromosomes carried duplications, all six sequenced chromosomes from the polytene division 18 hotspot carried small overlapping deletions within the same 110 kb intron of *kek5*. One possible explanation is that, owing to the size of *kek5*, conflict between replication and transcription generates a fragile site. Transcription through this long gene could delay replication, occasionally leaving this region unreplicated, possibly with an R-loop and single-stranded DNA present. This might generate a weak spot under tension at anaphase. CFS in humans have been shown to be enriched for long genes, and replication-transcription conflict is thought to contribute to their fragility^10,18^.

We found that breakpoints often occurred near preexisting transposon insertions, suggesting they may play a key role in determining the location of breakage events. It is especially notable that several termini coincide precisely with the locations of existing *FB4* DNA transposons, with DNA reads ending in *FB4* sequence (Figure 3B). *FB* (*foldback*) elements are known for their ability to form composite elements between nearby insertions of *FB* elements, and for their tendency to generate chromosome rearrangements^26^. This suggests a mechanism for this particular hotspot that involves mobilization by excision of a composite *FB4* element followed by mitosis. Formation of a dicentric would not be required since excision would leave a DSB. These events may have only been recovered here because of the strong selection for linearization of the ring^4^. It is also possible that the initial breaks did not occur at the *FB4* loci, but that the ends stabilized when they shortened to reach the *FB4* transposons.

Surprisingly, we find both DNA transposons and LTR retrotransposons near breakage sites. For DNA transposons, it is possible that their excision results in fragility. However, transposon excision cannot fully explain this pattern, as retrotransposons do not excise themselves during transposition. When envisioning how chromosomes break under tension, it would seem that stretching and unwinding of compacted chromatin must precede DNA breakage. It is possible that a unique chromatin state near transposons makes these regions more prone to unwinding and thus generate a breakage hotspot. However, little is known about the influence of local chromatin states on chromosome fragility under tension. Another possibility is that the unique sequence composition of transposons, rather than their transposition activity, makes these sites inherently fragile. This suggests that fragility is not tied to a specific class of transposon but may instead reflect general genomic features associated with transposons.

One pattern that unifies DNA transposons and retrotransposons is that they tend to be AT-rich due to codon bias^27,28^. AT-rich sequences are known to be more fragile in a variety of biological settings. For instance, in humans, CFS are known to be AT-rich^29^. At CFS, AT-rich regions often denature and form complex secondary structures that lead to replication fork stalling and underreplication^30^. It is possible that AT-rich retrotransposon sequences are a hotspot for replication fork stalling, which similarly leads to underreplication and fragility under tension during dicentric ring chromosome segregation. Additionally, we cannot rule out the possibility that the same chromosomal features that make some regions permissive to transposon insertions also make them prone to chromosome breakage under tension, which would generate a similar pattern of breaks at transposons.

An alternative possibility may involve tension-induced local denaturation of dsDNA at AT-rich sequences. Consistent with this idea, previous biophysical studies have shown that AT-rich sequences are more prone to denaturing into single strands under tension than GC-rich sequences^31,32^. Under this scenario, a fragile site is produced at AT-rich retrotransposons when local denaturation occurs. When only one strand of DNA bears the entirety of the tensile force, this region becomes the weakest link along the length of the chromosome, thus generating breakage hotspots. These hypotheses are not necessarily exclusive: replication-transcription conflict, replication fork stalling, and tension induced denaturation of DNA all lead to regions of ssDNA which could underly breakage hotspots. Regardless of the specific molecular mechanism, local stretches of single-stranded DNA, either pre-existing when cells enter mitosis or created by tension, may be the key determinant of dicentric breakage fragile sites.

## Data Availability Statement

Scripts can be found at https://github.com/JacksonRidges/dicentric_breakage. Raw sequencing data can be accessed through the SRA database with accession ID PRJNA1469658.

